# Thioesterase Superfamily Member 1 Undergoes Stimulus-coupled Reorganization to Regulate Metabolism

**DOI:** 10.1101/2020.02.18.954818

**Authors:** Yue Li, Norihiro Imai, Samaksh Goyal, Hayley T. Nicholls, Tibor I. Krisko, Mahnoor Baqai, Lay-Hong Ang, Matthew C. Tillman, Eric A. Ortlund, David E. Cohen, Susan J. Hagen

## Abstract

In brown adipose tissue, cold exposure promotes thermogenesis, in large part by increasing mitochondrial β-oxidation of lipid droplet-derived fatty acids. This process is suppressed by thioesterase superfamily member 1 (Them1), a long chain fatty acyl-CoA thioesterase that is highly upregulated by cold ambient temperatures. Them1 reduces fatty acid availability for β-oxidation in mitochondria and limits thermogenesis by cellular mechanisms that are not well defined. We show that Them1 regulates metabolism by undergoing marked intracellular conformational changes that occur in response to β-adrenergic stimulation. Mechanistically, Them1 formed puncta that were localized near LD and mitochondria in an immortalized brown adipose cell line. In response to stimulation by norepinephrine, Them1 was phosphorylated by PKCβ at S15, which specifically inhibited puncta formation and resulted in a diffuse intracellular localization. This change in Them1 localization also occurred after stimulation in vivo. Puncta formation activated Them1 metabolic activity in vitro, as evidenced by suppression of oxygen consumption following β-adrenergic stimulation. We show by correlative light and electron microscopy that puncta are biomolecular condensates (also known as membraneless organelles) that typically form by phase separation. Them1 contains one intrinsically disordered region at the N-terminus with multiple interacting motifs that is frequently observed in phase-separating proteins. Phosphorylation, which is known to disrupt phase separation and aggregation, results in a diffuse Them1 localization. Our data thus establish that Them1 forms intracellular biomolecular condensates that limit fatty acid oxidation and suppress thermogenesis. During a period of energy demand, the condensates are disrupted by phosphorylation to allow for maximal thermogenesis. The stimulus-coupled reorganization of Them1 thus provides fine-tuning of thermogenesis and energy expenditure.

## Introduction

Intracellular triglycerides are stored within lipid droplets (LD) that are juxtaposed to mitochondria in the cytoplasm^1^. This close relationship facilitates the rapid transfer of fatty acids to mitochondria, which are generated by lipolysis in response to cellular energy demands^2^. One such demand is non-shivering thermogenesis, which is mediated by brown adipose tissue (BAT) in response to cold exposure to generate heat^3^. This occurs, at least in part, when mitochondrial β-oxidation is uncoupled from oxidative phosphorylation via uncoupling protein 1. The hydrolysis of LD-derived triglycerides is initiated following the release of norepinephrine from neurons, which binds to β3 adrenergic receptors on brown adipocytes to stimulate a cell signaling cascade that activates adenylyl cyclase to generate cAMP and activate protein kinase A (PKA)^4^. In brown adipocytes, the generation of fatty acids for mitochondrial oxidation requires PKA to phosphorylate/activate perilipin, a regulatory membrane-bound protein that encircles LD. The function of perilipin is to protect triglycerides within the LD from lipolysis, and its phosphorylation by PKA allows access of cytoplasmic adipose triglyceride lipase (ATGL), hormone sensitive lipase (HSL), and monoacylglycerol lipase (MAGL), to generate free fatty acids through sequential lipolytic steps^5^. Once fatty acids are free in the cytoplasm, they are esterified with coenzyme A (CoA) by long chain acyl-CoA synthetase 1 (ACSL1) to form fatty acyl-CoAs, which are transported into mitochondria via carnitine palmitoyl transferase 1 and metabolized to produce heat^6^. Fatty acyl-CoA molecules can also be hydrolyzed to fatty acids in the cytoplasm by acyl-CoA-thioesterase (Acot) isoforms^7,8^. The fatty acids generated can be utilized to make additional fatty acyl-CoAs that are transported into mitochondria or can be returned to the LD for storage.

Thioesterase superfamily member 1 (Them1), which is also known as brown fat inducible thioesterase or steroidogenic acute regulatory protein-related lipid transfer (START) domain 14 (StarD14)/Acot11, plays an important role in energy homeostasis^9,10^. Mice with the genetic deletion of Them1 exhibit increased energy expenditure, which results in reduced weight gain when challenged with a high fat diet despite high food consumption^10^. This is attributable to increased mitochondrial fatty acid oxidation, and mechanistic studies have demonstrated that Them1 functions to suppress energy expenditure by limiting triglyceride hydrolysis in BAT, thus inhibiting the mitochondrial oxidation of LD-derived fatty acids^10,11^.

Them1 is comprised of tandem N-terminal thioesterase domains and a C-terminal lipid-binding domain steroidogenic acute regulatory-related lipid transfer (START) domain. Proteomics has revealed that there are three BAT-selective phosphorylation sites near the N-terminus of Them1 at serine (S) 15, 18, and 25^12^. Based on an *in silico* analysis suggesting that protein kinase C β (PKCβ) may be responsible for phosphorylation at these sites^13,14^, we explored the mechanistic contribution of PKCβ-mediated Them1 phosphorylation to the cellular regulation of fatty acid oxidation in BAT. Here, we show that Them1 exists in two different phosphorylation-dependent conformational states: punctate and diffuse. Punctate refers to small, highly concentrated aggregations of Them1 that localize to regions of the cytoplasm that interface with LD and mitochondria, whereas diffuse refers to homogeneously distributed Them1 across the cell cytoplasm and nucleus. Transformation of these conformational states occurs by the redistribution of existing Them1 and not by new protein synthesis. We demonstrate that β-adrenergic stimulation leads to the activation of PKCβ downstream of PKA, leading to site specific phosphorylation of Them1 at S15. This drives the dissolution of puncta both in vitro and in vivo. A functional analysis revealed that Them1 in punctate form suppresses fatty acid oxidation, whereas this suppression is abrogated when it is diffusely distributed in the cell cytoplasm. Overall, these findings highlight the importance of Them1 phosphorylation in the regulation of thermogenesis in BAT and lend support for targeting Them1 for the management of obesity-related disorders.

## Results

### Identification of PKCβ as the putative kinase for N-terminal serine phosphorylation of Them1

To assess the BAT-selective phosphorylation of Them1, we consulted the phosphomouse database (http://gygi.med.harvard.edu/phosphomouse), which revealed phosphopeptides containing phosphorylation events at S15, S18 and S25. Interestingly, Them1 S15 and S25 are highly conserved; S18 is also conserved but less so because the serine residue can be substituted with alanine (Supplementary Fig. 1). PKCβ is a predicted kinase for these sites^13,14^ and like Them1, PKCβ activity limits energy expenditure^15^ and promotes diet-induced obesity and non-alcoholic fatty liver disease^16,17^.

### Phosphorylation-dependent changes in Them1 localization after stimulation

Cultured immortalized brown adipose (iBAs) cells did not express Them1 mRNA or protein (Supplementary Fig. 2a), as was the case for primary cultured brown adipocytes^11^. Instead, Them1 expression required transfection to study its biochemical and physiological characteristics in vitro. To determine the intracellular localization of Them1, we first transfected differentiated iBAs cells with Them1 (full-length)-EGFP and then analyzed Them1 expression in fixed or living cells by confocal microscopy (Fig. 1). Them1 exhibited a punctate intracellular localization (Fig. 1a). When cells co-stained for LD and mitochondria were reconstructed from 3-D stacks, Them1 was closely associated, but did not co-localize, with either of these organelles (Fig. 1b). Aggregation of Them1 was not due to EGFP, as evidenced by the diffuse intracellular localization of EGFP in cells transfected with EGFP alone (Supplementary Fig. 3a). Furthermore, Them1 formed puncta in transfected cells without the EFGP tag (Supplementary Fig. 3b).

**Fig. 1.**
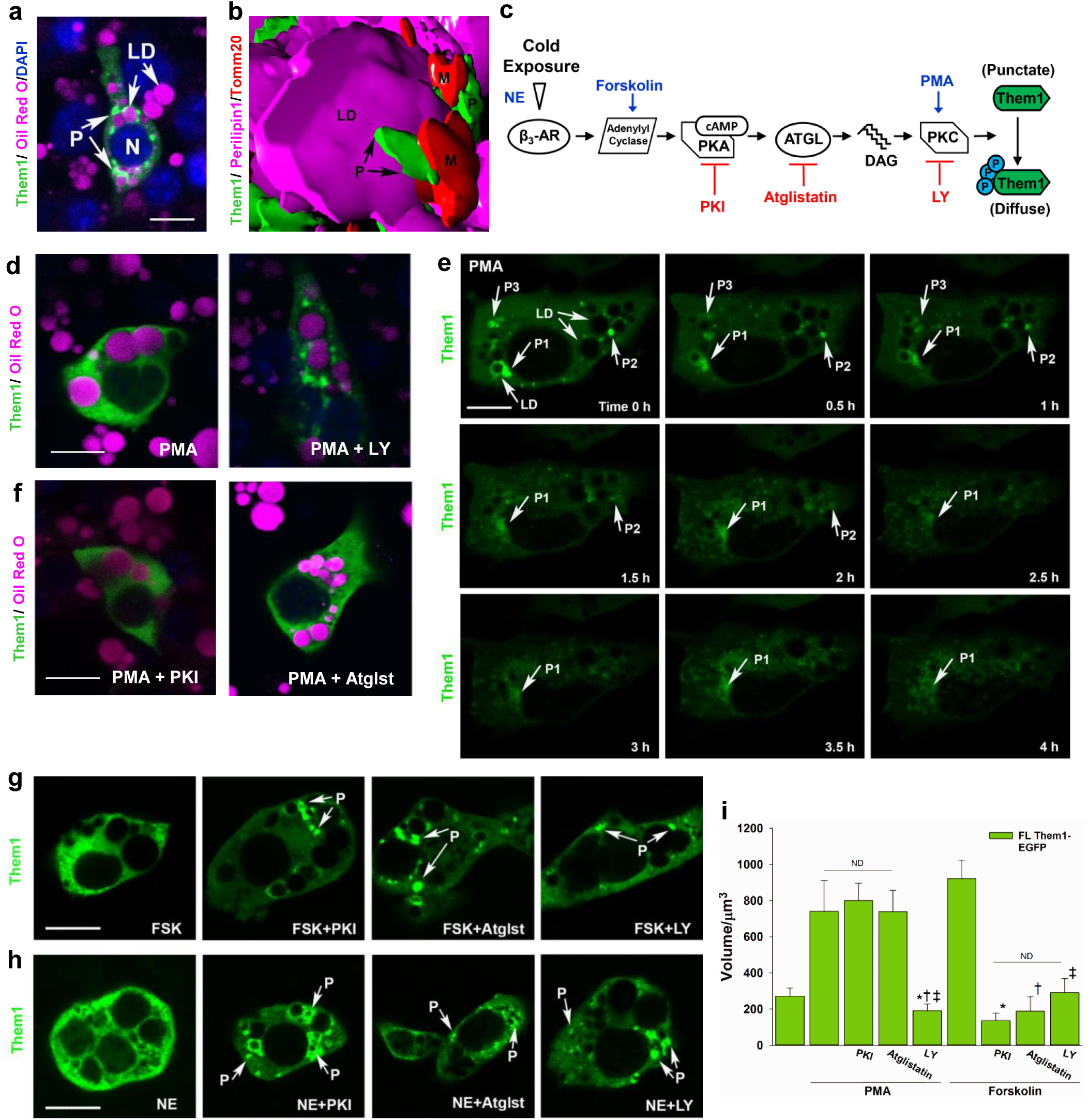
Regulation of Them1 subcellular localization in immortalized brown adipocytes (iBAs) by PKCβ. **a,** Confocal microscopy of iBAs expressing Them1-EGFP (green). Them1 forms puncta (P) in non-stimulated cells, which localize near lipid droplets (LD). N, Nucleus. **b,** 3-D image of a iBA expressing Them1-EGFP, perilipin1 to identify lipid droplets (LD), and Tomm20 to identify mitochondria (M). This image was processed using Volocity software and is shown in surface projection mode to demonstrate that Them1-containing puncta (P) localize between LD and mitochondria. **c,** Schematic illustration of the putative Them1 regulatory pathway required for the phosphorylation and 3-D organization of Them1. **d**, PMA-induced activation of iBAs expressing Them1-EGFP results in the intracellular diffusion of Them1, which can be blocked by the PKCβ inhibitor LY333531 (LY). **e,** Live-cell imaging was used to record changes in Them1 localization over time after PMA treatment. The initial punctate spots at T0 (P1, P2, P3) redistribute across the cytoplasm over time and by 4 h after stimulation are not present in the cytoplasm. LD, lipid droplets. **f,** After PMA-induced PKC activation (see panel c), neither PKI or atglistatin (Atglst) blocked the diffusion of Them1 from puncta. **g,** Whereas the activation of PKA and PKC by forskolin (FSK) resulted in the diffusion of Them1, PKI, atglistatin (Atglst), and LY333531 (LY) blocked the diffusion of puncta (P). **h,** Activation of the β3 adrenergic receptor by norepinephrine (NE), which mimics cold exposure in BAT cells (see panel c), causes the diffusion of Them1 from puncta. However, NE treatment in presence of PKI, atgilstatin (Atgilst), or LY333531 (LY) blocked the diffusion of puncta (P). Scale bars in a-h, 10 μm. **i,** Quantification of results from PMA or forskolin stimulation with or without pathway inhibitors PKI, atgilstatin, or LY333531. When Them1 is in puncta, it is concentrated in a small intracellular volume. In contrast, diffuse Them1 occupies a large cytoplasmic volume. Bars represent the mean +/− s.e.m and *n* is from the following number of images from 3 different experiments: Them1-EGFP, *n* = 17; PMA, *n* = 8; PMA+PKI, *n* = 16; PMA+Atglst, *n* = 7; PMA+LY333531, *n* = 12; FSK, *n* = 9; FSK+PKI, *n* = 12; FSK+Atglst, *n* = 7; FSK+LY333531, *n* = 10. Three to five cells per independent experiment were evaluated. *** *P*< 0.001 and ND represents no statistical significance as determined by a one-way ANOVA.

We next explored the hypothesis that the stimulus-mediated phosphorylation of Them1 facilitates changes in its localization (Fig. 1c). For this, differentiated iBAs cells were incubated with phorbol 12-myristate 13-acetate (PMA), and the localization of Them1 was examined in fixed or living cells (Fig. 1d,e). PMA was used to activate PKC (Fig. 1c), which initiated the time-dependent dissolution of Them1 from puncta (Fig. 1d,e). The dissolution of puncta resulted in Them1 uniformly distributed within the cell cytoplasm (Fig. 1d), which occurred over a 4 h period as determined using time-lapse live cell microscopy (Fig. 1e). Consistent with a role for PKCβ in Them1 phosphorylation, the isoform-specific PKCβ inhibitor ruboxistaurin (LY333531) completely blocked the dissolution of puncta in PMA-treated cells (Fig. 1d).

We next sought to identify upstream signaling events and define the pathway leading to PKCβ-mediated Them1 phosphorylation (Fig. 1c). iBAs cells incubated with the ATGL inhibitor atglistatin or the PKA inhibitor PKI prior to stimulation with PMA had a diffuse Them1 distribution in the cytoplasm (Fig. 1f). Similarly, iBAs cells stimulated with forskolin (Fig. 1g), which activates adenyl cyclase and PKA upstream of PKC (Fig. 1c), or with norepinephrine (Fig. 1h), which activates both PKA and PKC (Fig. 1c) resulted in diffusely localized Them1, which was blocked by inhibiting downstream effectors with PKI, Agistatin, or LY, respectively (Fig. 1g, h). Measuring the volume of intracellular Them1 in iBAs cells transfected with Them1-EGFP further demonstrated that Them1 in puncta is concentrated and occupies significantly less volume than occurs when Them1-EGFP is diffusely distributed after stimulation with PMA or forskolin or when the phosphorylation by PKCβ is inhibited (Fig. 1i). Additionally, the dissolution of Them1 from puncta after stimulation with PMA or forskolin was not attributable to the synthesis of new Them1-EGFP protein, because inhibition of protein synthesis using cyclohexamide during the 4 h period of stimulation had no effect on Them1 localization or protein concentration (Supplementary Fig. 3). Overall, these results suggest that norepinephrine stimulation after cold exposure activates PKA, and the activation of PKC occurs downstream of PKA activation (Fig. 1c). Furthermore, they identify PKCβ-mediated phosphorylation of Them1 as being responsible for the relocalization of Them1 after stimulation (Fig. 1c).

We next sought evidence that the stimulus-coupled changes in Them1 localization also occurred in brown adipose cells in vivo (Fig. 2). For this, mice were injected with saline (control) or with the β3 adrenergic receptor selective agonist CL316,243. In saline-injected mice at baseline, Them1 formed numerous distinct puncta near LD and in the cytoplasm (Fig. 2a,c). By 2 and 4 h after saline injection, however, the volume of Them1 and the number of puncta was reduced (Fig. 2a,c). The high Them1 at baseline was due to 24 h of mild cold exposure that increases Them1 in BAT (see Methods), which was reduced by returning mice to thermoneutrality and a stress-effect caused by saline injections alone.^18^

**Fig. 2.**
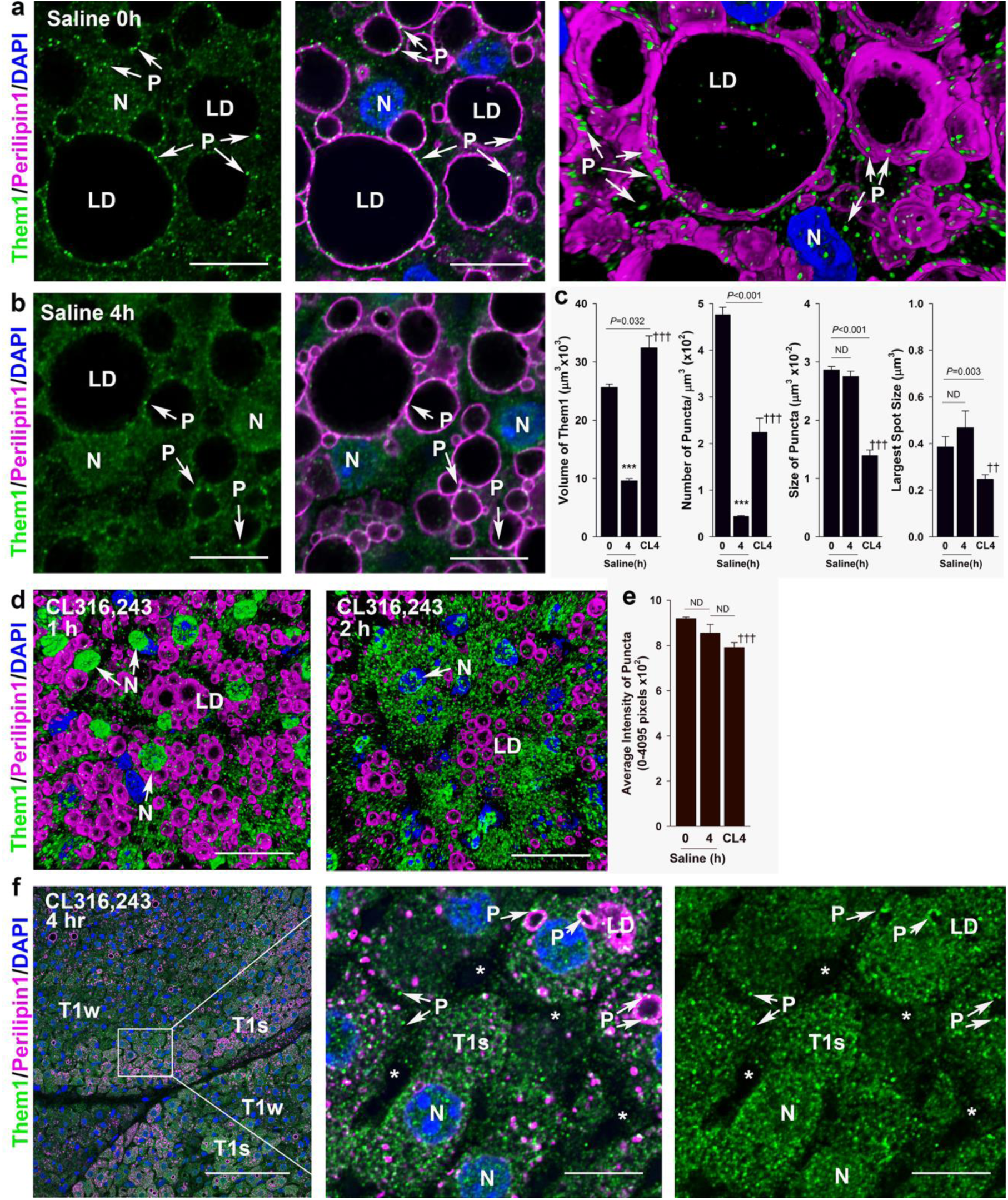
Them1-containing puncta in brown adipose tissue (BAT) diffuse after stimulation in vivo. **a**, BAT was excised from mice immediately after the administration of saline (baseline). These tissues consisted of large lipid droplets (LD) surrounded by perilipin and numerous Them1-containing puncta (P) that were localized in the cytoplasm or in association with the outer edge of perilipin at the lateral edge of LD. N, nucleus. Scale bar, 10 μm. **b**, BAT tissues from mice injected with saline and then tissues collected 4 h later. Them1 expression was lower and the number of puncta reduced. N, nucleus. Scale bar, 10 μm. **c**, Quantification of the 3-D volume of Them1, the number of puncta/area, the size of puncta, and the size of the largest puncta in mice injected with saline (control) or CL316,243, a β3 adrenergic receptor agonist. Error bars represent mean + s.e.m. from n=9 images at saline 0h and 18 images at saline 4h and CL 4h. *P* values were determined using the Student’s t-test. ***, significantly different at *P*<0.001 compared to saline 0h, †††, significantly different at *P*<0.001 compared to saline 4h. **d**, BAT tissues from mice exposed to CL316,243 for 1 h. In these tissues, lipid droplets (LD) were smaller than in saline-treated mice and the majority of Them1 was associated with the outer edge of nuclei (N). By 2 h after CL316,243 administration, LD were smaller than in BAT tissues after 1 h of CL exposure and Them1 occupied a large cell volume at this time point. **e**, Quantification of the average intensity of puncta in mice injected with saline (control) compared to CL316,243. Error bars represent mean + s.e.m. from n=9 images at saline 0h and 18 images at saline 4h and CL 4h. *P* values were determined using the Student’s t-test. †††, significantly different at *P*<0.001 compared to saline 0h. **f**, BAT tissue from mice 4 h after the administration of CL316,243. LD were small and Them1 was found in areas with weak fluorescence signal (T1w) or with strong fluorescence signal (T1s). In areas with strong signal (higher mag images) Them1 was in puncta (P) that were weaker and smaller than found in saline control tissues. N, Nucleus. *Increased space between BAT cells due to the lack of lipid droplets and thus smaller cell size. Scale bars, 10 μm.

By 1 h after CL316,243 administration, LD were reduced in size and number due to lipid hydrolysis and Them1 aggregated at the outer nuclear membrane in most cells (Fig. 2d). By 2 h after CL administration, LD were smaller yet and Them1 puncta occupied a large cytoplasmic volume (Fig. 2d). By 4 h after CL administration, there were few intracellular LD as evidenced by the loss and disruption of perilipin 1 expression, and Them1 occupied a large cytoplasmic volume (Fig. 2c,f). Interestingly, Them1 localization was strong in some cells and weak in others (Fig. 2f), possibly reflecting differences in the metabolic activity of different cell populations in BAT.^19^ Although punctate-looking structures remained in the cytoplasm, the number of puncta, size of puncta, largest spot size of puncta, and the average intensity of puncta were reduced compared to cells from saline-injected mice at baseline (Fig. 2c,e,f). When taken together with observations in vitro using iBAs cells, these in vivo studies support the notion that reorganization of Them1 puncta occurs after stimulation with norepinephrine.

### Phosphorylation of Them1 at S15 drives changes in Them1 localization after stimulation

To develop direct evidence that residues S15, S18, and S25 of Them1 are involved in puncta formation and dissolution, we deleted the first 36 aa of Them1 to generate a construct (Δ1-36-EGFP) starting at methionine (M)37 that did not contain these 3 phosphorylation sites (Fig. 3a, b). The Δ1-36 mutant of Them1 exhibited a diffuse localization (Fig. 3c). We then linked a peptide containing the first 36 aa of Them1 directly to EGFP (not shown), which exhibited the same punctate localization as full length Them1 (Fig. 3c). These results strongly support that amino acids 1-36 direct Them1 localization to puncta and suggest that phosphorylation of Them1 at one or more serine residues in the N-terminal region drives the conformational change in Them1 after stimulation.

**Fig. 3.**
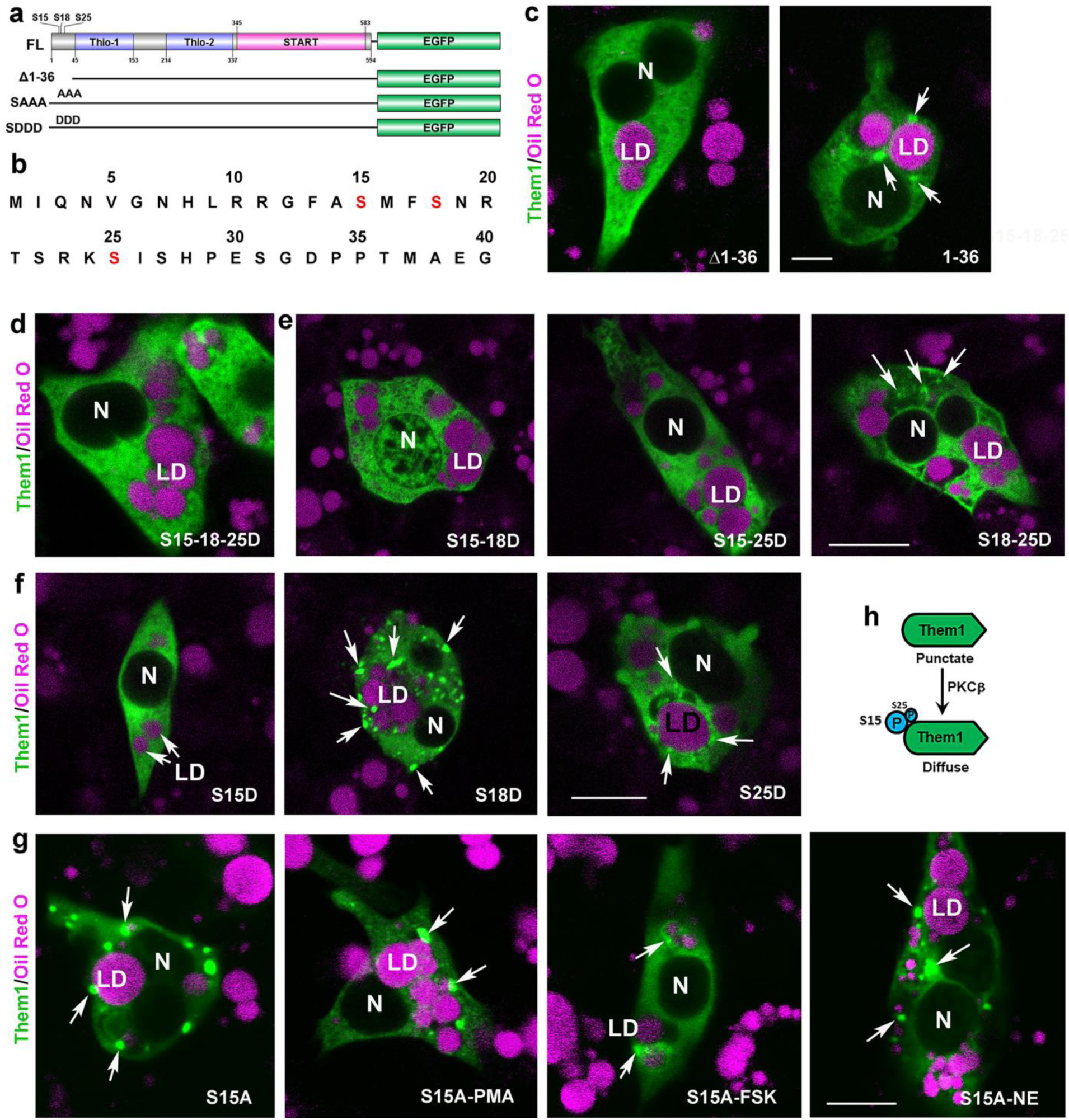
Phosphorylation at S15 in the N-terminus regulates the diffusion of Them1 in iBAs. **a, b** Illustration of the Them1 structure and its mutant constructs tagged with EGFP used in the experiments. FL, full-length Them1 linked to EGFP at the carboxy terminus. Δ1-36, deletion of amino acids 1-36 in the Them1 sequence; 1-36, a construct containing only the first 36 amino acids in the Them1 sequence (not shown); AAA, serine residues at amino acids 15, 18 and 25 were mutated to alanine; DDD, serine residues at amino acids 15, 18 and 25 were mutated to aspartic acid. All mutant Them1 constructs were linked to EGFP. **c,** the Δ1-36 Them1 construct diffuses in the cytoplasm whereas the 1-36 construct forms puncta around lipid droplets (LD). **d,** Mutation of S15/S18/S25 to aspartic acid (DDD) results in diffuse Them1 localization. **e**, S15/S18 or S15/S25 mutant constructs result in diffuse Them1 localization whereas the S18/S25 construct expressed some puncta and some diffuse Them1. **f,** The single mutation at S15, and to some extent S25, but not S18 to aspartic acid resulted in the diffusion of Them1. **g,** The serine to alanine mutation at S15 results in Them1-containing puncta, which are not dispersed after activation by PMA, FSK or NE. Scale bar in c-g, 10 μm. For each construct, the data are representative of *n* = 3 independent experiments with 3-5 individual cells photographed per treatment. **h**, Schematic diagram of Them1 localization before and after PKCβ-mediated phosphorylation of S15, and to a lesser degree S25, at the N-terminus.

To further explore this idea, we mutated each putative phosphorylation site by exchanging S for aspartic acid (S15-18-25D, or DDD), which mimicked the phosphorylated state of the S amino acid residues (Fig. 3a). This construct resulted in diffuse Them1 localization (Fig. 3d). When the S sites were mutated in pairs (Fig. 3e), or individually (Fig. 3f), the combinations including S15D, and to a minor extent S25, led to diffuse Them1 localization (Fig. 3e, f). By contrast, exchanging S for alanine at S15, which is unable to be phosphorylated, resulted in the punctate localization of Them1 alone or after stimulation with PMA, FSK, or norepinephrine (Fig. 3g). Overall, these results demonstrate that the phosphorylation of Them1 at highly conserved S residues, S15 with a minor contribution from S25 (Fig. 3h), is both necessary and sufficient to result in the diffuse localization of Them1 in iBAs cells.

### Puncta formed by Them1 are metabolically active structures

Although plasmid-mediated transduction of Them1 constructs was sufficient for fluorescence-based imaging studies, the low transfection efficiencies prevented quantitative assessments of the impact of puncta formation on cellular metabolism. To obtain higher transfection efficiency, we constructed adenoviral vectors. These included wild type Them1 or Them1 with AAA or DDD substitutions and EGFP fused at the carboxy terminus (Fig. 3a, b). An EGFP-expressing adenoviral vector was used as a control. Preliminary experiments established that an MOI of 1:40 yielded Them1 expression that was similar to the tissue expression level of the protein in BAT of cold-exposed mice (Supplementary Fig. 2b) with a transfection efficiency of approximately 45%. Both wild type (not shown) and AAA (Fig. 4a) substituted Ad-Them1-EGFP formed puncta in iBAs cells. Using near super-resolution imaging via Zeiss Airyscan, it was clear from 3-D images that Them1 resides in discrete puncta that were closely associated with LD (Fig. 4a). This localization was consistent with the results from our plasmid constructs. In contrast, DDD substituted Them1 exhibited a diffuse localization (Fig. 4b). In the near super-resolution 3-D images, diffuse Them1 filled the entire volume of cytoplasm not occupied by cellular organelles (Fig. 4b) consistent with the results from our plasmid constructs.

**Fig. 4.**
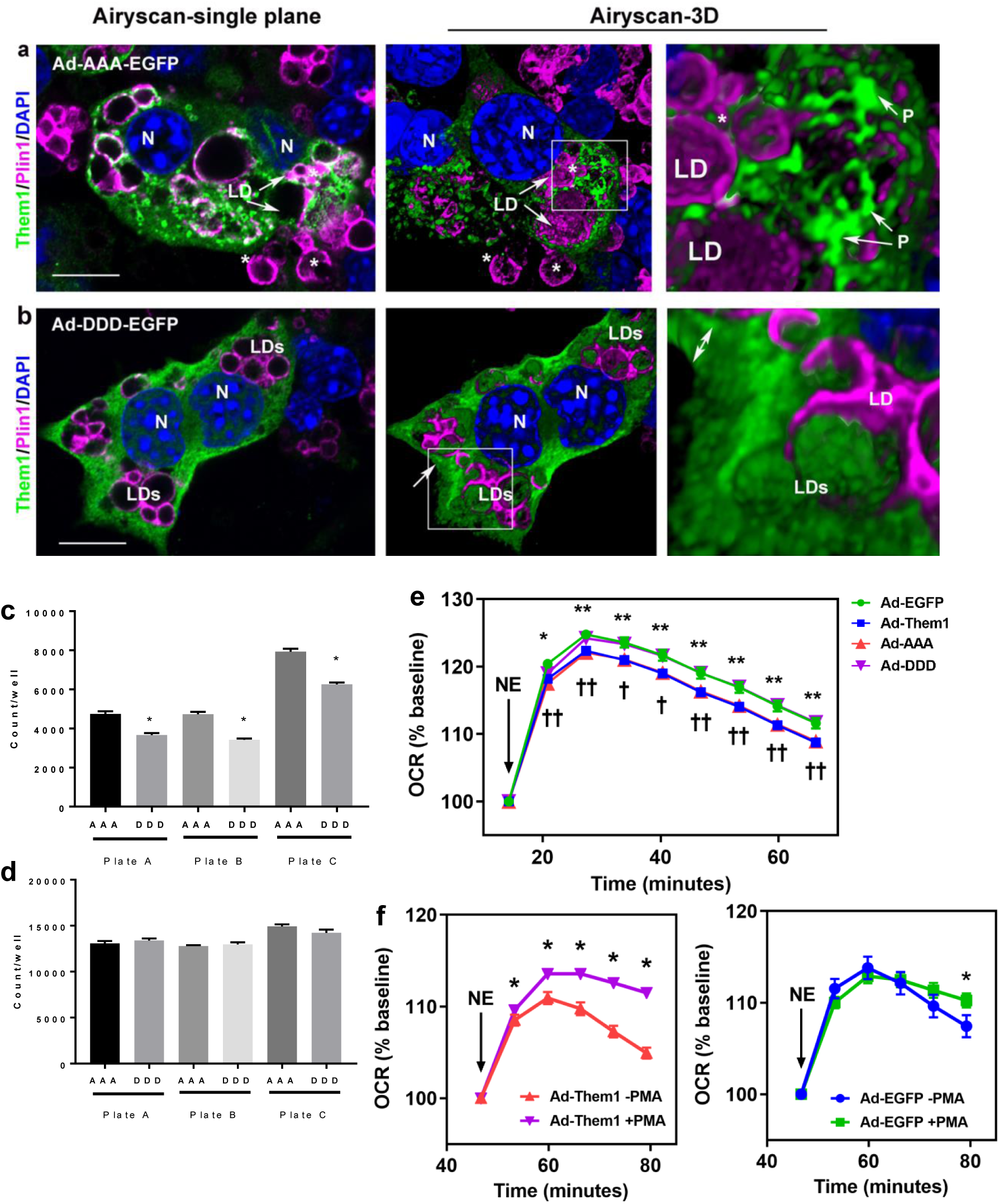
Adenoviral transduction of iBAs with full-length Them1 and mutant constructs confirms that the punctate localization of Them1 is metabolically active. **a,** adenovirus-mediated expression of Them1-EGFP with S15, S18 and S25 mutated to alanine (Av-AAA) formed puncta near lipid droplets (LD). This close relationship was highlighted by using near super-resolution Zeiss Airyscan imaging and reconstructing cells in 3-D. Serine to aspartic acid mutation at S15, S18 and S25 (Av-DDD) caused diffusion of Them1-EGFP. In 3-D near super-resolution Zeiss Airyscan images, Them1 filled the thickness of the cytoplasm (double white arrow) and could be found above and below the plane of organelles including lipid droplets (LD), which excluded Them1. N, nucleus. Scale bars, 10 μm. Data are representative of *n* = 3 independent experiments. **c**, Calculation of the total number of EGFP-positive cells/plate, which was used to normalize metabolic data in **e** and **f**. **d**, Calculation of the total number of living cells/plate. These data demonstrate that Ad-AAA or Ad-DDD transduction does not change the number of viable cells. **e**, iBAs were transduced with Ad-full length Them1 (FL-Them1), Ad-Them1(AAA), Ad-Them1(DDD) and Ad-EGFP (control) and treated with 3 μM phorbol 12-myristate 13-acetate (PMA) thirty minutes before norepinephrine (NE) stimulation. The response of OCR values to stimulation with 1 μM NE was measured as % of baseline for 60 min and normalized to the total number of EGFP-positive cells. The graph shows combined data for *n* = 3 independent experiments. Error bars indicate mean +/− s.e.m; EGFP vs FL-Them1 *P<0.05, **P<0.01 and AAA vs DDD † P<0.05, †† P<0.01 (Student’s t-test). **f**, Ad-Them1 or Ad-EGFP reconstituted iBAs cells with or without 3 μM phorbol 12-myristate 13-acetate (PMA) thirty minutes before NE stimulation. Relative OCR values were calculated based on baseline OCR values before NE injection, as was done in panel **e**. Graph shows combined data from *n* = 3 independent experiments. *P<0.05 (Student’s t-test).

We next examined the role of Them1 in regulating the oxygen consumption rate (OCR) in differentiated iBAs cells in response to norepinephrine stimulation. OCR values are a surrogate measure of oxidation rates of fatty acids generated by hydrolysis of LD-derived triglycerides^11^. After optimizing conditions of cell density by calculating the total number of EGFP-containing cells/total cells per plate (Fig. 4c,d), OCR values were measured at baseline and after norepinephrine stimulation. Baseline and norepinephrine-stimulated OCR values were similar for Ad-DDD-EGFP and the Ad-EGFP control, whereas norepinephrine-stimulated values of OCR for Ad-Them1 and Ad-AAA-EGFP were reduced (Fig. 4e). Because the stimulated redistribution of Ad-Them1 to become fully diffuse occurs over a period of 4 h (Fig. 1d), increases in OCR values within the first 1 h after norepinephrine exposure (Fig. 4e) would be expected to reflect only partial dissolution of Them1-containing puncta. By pre-incubating iBAs with PMA prior to norepinephrine exposure, which would result in the further dissolution of puncta, we observe an additional increase in OCR values relative to no PMA, whereas no effects are observed using Ad-EGFP controls (Fig. 4f). These findings indicate that Them1 is active in suppressing LD-derived FA oxidation when in its punctate configuration.

### Puncta are biomolecular condensates, or “membraneless organelles”, by correlative light/electron microscopy

To examine the ultrastructural details of puncta in iBAs cells, we next used the Them1-EGFP vector constructs that included an ascorbate peroxidase–derived APEX2 tag appended to the carboxy-terminus to perform correlative light/electron microscopy (Fig. 5). The APEX2 tag was developed with diaminobenzadine, resulting in brown reaction product only in cells expressing Them1 (Fig. 5a). This procedure allowed us to clearly determine which cells were transfected and expressing Them1. Them1 puncta were imaged using the AAA vector construct (Fig. 5b,c), and diffuse Them1 was evaluated using the DDD vector construct (Fig. 5d). Cells transfected with EGFP-APEX2 without Them1 showed pale staining uniformly throughout cells including the nucleus.

**Fig. 5.**
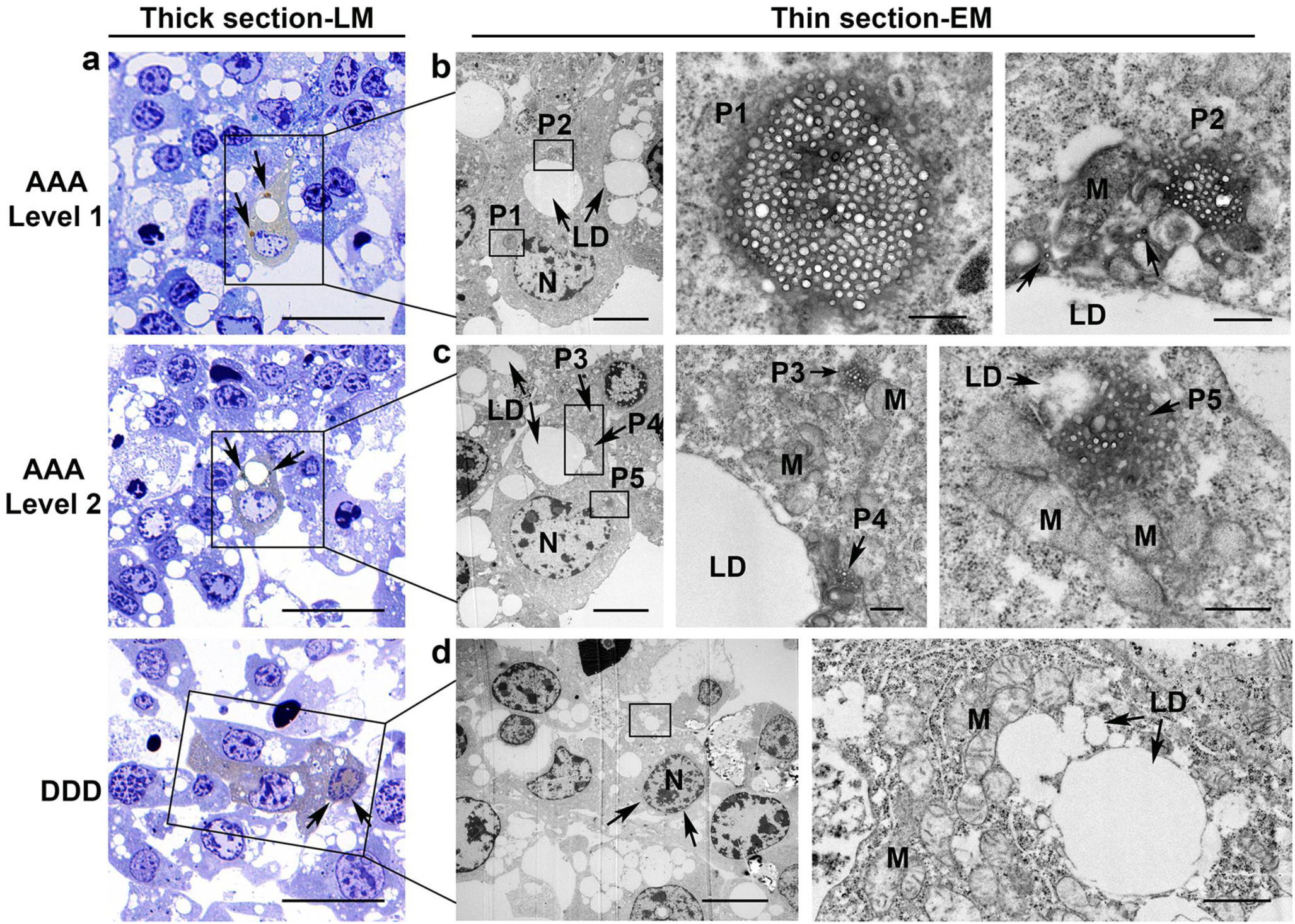
Correlative light and electron microscopy identifies puncta as biomolecular condensates. **a,** Thick, 0.5 μm, plastic sections of iBAs in culture transfected with phosphorylation mutants AAA- or DDD-Them1-EGFP/APEX2. Cells stained brown (box) also have dark brown punctate structures in mutant AAA-Them1 expressing cells (arrows). For AAA-transfected cells, serial sections show two different levels of the same cell. No puncta are present in cells transfected with the DDD-Them1 mutant although Them1 can also be found in the nucleus (arrows). Scale bar, 25 μm. **b**, In thin sections of the boxed cell in **a**, puncta 1 (P1) is localized near the nucleus (N) and puncta 2 (P2) is localized near a lipid droplet (LD). At high magnification, punctate structures, P1 and P2, have no surrounding membrane and consist of liquid droplets embedded in an amorphous matrix. Scale bars, 5 μm (low magnification) and 0.5 μm (high magnification). **c**, In thin sections of the boxed cell in **a**, this section shows smaller puncta in the cytoplasm (P3), puncta associated with lipid droplets (LD; P4), and puncta in close association with LD and mitochondria (M) in the cell cytoplasm (P5). Scale bars, 5 μm (low magnification) and 0.5 μm (high magnification). **d**, In thin sections from the boxed region in **a**, no puncta are found in cells transfected with the DDD-Them1 mutant. Them1 is distributed throughout the cytoplasm and in the nucleus (N; arrows). Scale bars, 10 μm (low magnification) and 1 μm (higher magnification). All images are representative of the results obtained from *n* = 3 independent experiments.

Consistent with fluorescence micrographs of iBAs cells labeled with Them1-EGFP (Fig. 1a), puncta in Them1-EGFP-APEX2 cells were found exclusively in the cytoplasm in close proximity to both LD and mitochondria (Fig. 5a-c). Although the puncta resembled mitochondria from steroid-secreting cells^20^, there was no apparent membrane surrounding the structure. Instead, puncta were an aggregate of small round and elongate droplets with amorphous boundaries (Fig. 5b,c). Diffusely localized Them1 was not clearly associated with any cytoplasmic structure (Fig. 5d).

Because puncta were membraneless structures with characteristics similar to that described for biomolecular condensates within the cell cytoplasm^21^, we evaluated the Them1 primary sequence for characteristics that would facilitate a phase separation, aggregation, and/or the formation of puncta (Fig. 6). Bioinformatic analysis of the amino acid sequence for Them1 (Fig. 6a) showed a highly disordered region at the N-terminus spanning residues 20-45 as predicted by IUPRED (prediction of intrinsic disorder) curves (Fig. 6b). There were no prion-like regions in the sequence as determined by PLD (Fig. 6c) or FOLD curves (Supplementary Fig. 4b). The N-terminus contains a high proportion of charged residues with a patch of basic residues followed by a patch of acidic residues (Supplementary Fig. 4c,d), which could engage in non-covalent crosslinking with other proteins or itself to drive a phase transition^22^. However, the PScore, a pi-pi interaction predictor, did not reach significance (Fig. 6d), which suggests that the sequence of Them1 does not have the propensity to phase separate based on pi-pi interactions. ANCHOR2 analysis, which predicts protein binding regions in disordered proteins, showed that the N-terminal 20 amino acids were a highly disordered binding region (Fig. 6e). Interestingly, two of the phosphorylation sites, S15 and S18, lie in this region (Fig. 6a), suggesting that the phosphorylation sites may regulate binding events. Because multivalent interactions between proteins can lead to a phase separation^21^, we also performed a motif scan on the first 20 amino acids. This analysis suggested a weak association of these amino acids with a formin homology-2 (FH2) domain^23^, which promotes the growth of unbranched actin filaments.

**Fig. 6.**
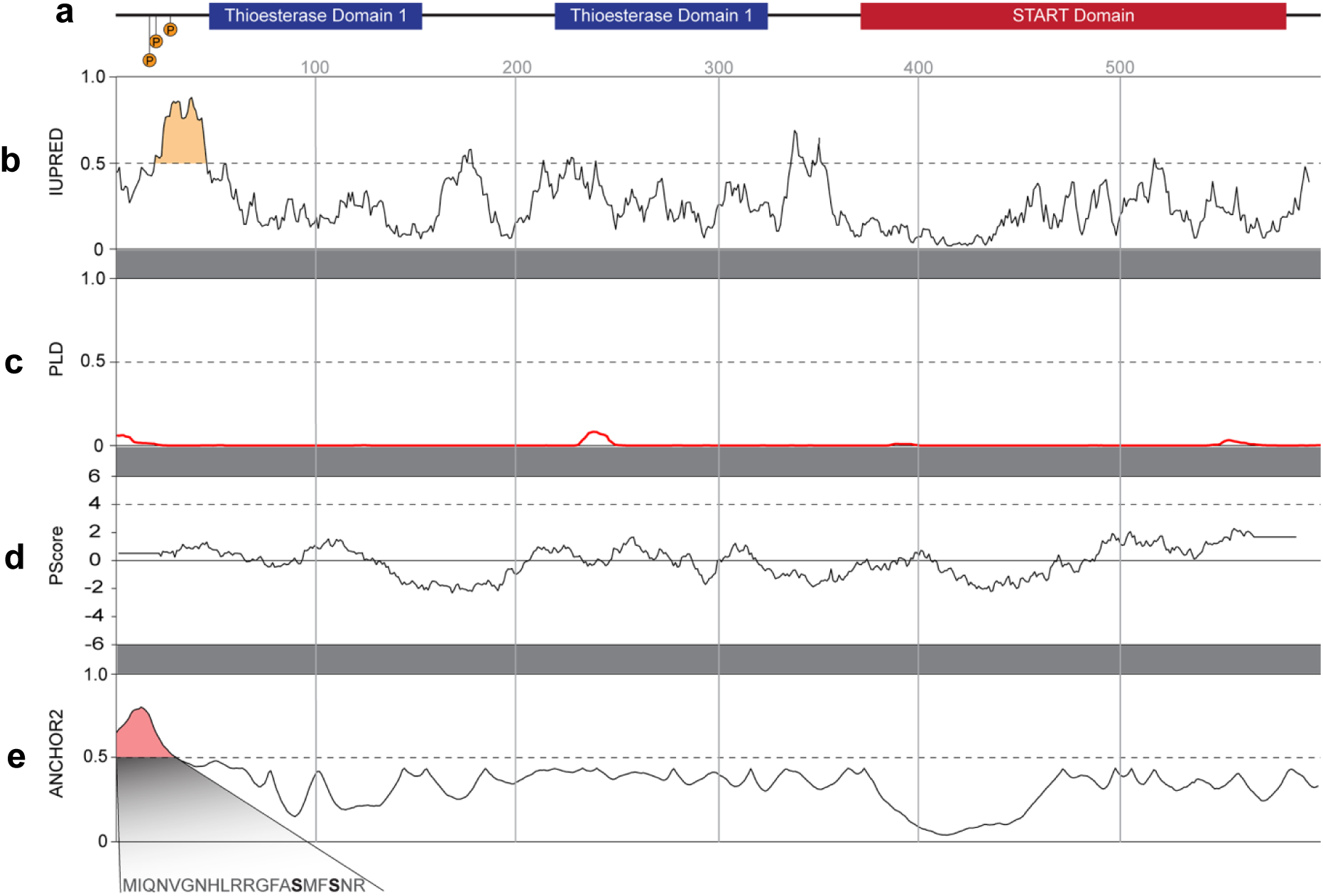
Bioinformatic analysis of Them1 suggests that a disordered region at the N-terminus spanning residues 20-45 may be involved in puncta formation. **a**, Schematic diagram of the Them1 amino acid sequence aligned with the amino acid number from the N- to C-terminus. **b**, Results from the IUPRED server, which predicts regions of disorder. The dotted line signifies the threshold for significance. The data indicate a large region of disorder in the first 50 amino acids (highlighted in gold). **c**, Results from the Prion-like Amino Acid Composition server. The dotted line at 0.5 is a threshold for significance. The graph, red line, demonstrates no prion-like regions in the Them1 amino acid sequence. **d**, Results from the PScore, which is a predictor of the propensity to phase separate based on pi-pi interactions. The dotted line is a threshold set at 4 to determine significance. The amino acid sequence of Them1 did not reach significance. **e**, Results from an ANCHOR2 analysis, which predicts protein binding regions in disordered proteins. This analysis demonstrates that the N-terminal 20 amino acids of Them1 were strongly predicted to be a disordered binding region (highlighted in red). Two of the phosphorylation sites lie in this region (expanded region, also refer to Fig. 3b), suggesting they could potentially regulate a binding event.

## Discussion

The expression and functional impact of Them1 in thermogenesis is unique. Them1 was first identified as a gene that is markedly upregulated in response to cold ambient temperatures and suppressed by warm temperatures^9^. Whereas this initially suggested that Them1 function was to promote thermogenesis^9^, homozygous disruption of *Them1* in mice led to increased energy expenditure^10,11^. This result indicated that Them1 functions, on balance, to suppress energy expenditure. The current study helps to explain how a suppressor of energy expenditure can be induced by cold ambient temperatures, which also upregulates thermogenic genes; Them1 is phosphorylated and inactivated by the same adrenergic stimulus that promotes thermogenesis. Additionally, our studies have demonstrated phosphorylation-dependent intracellular redistribution of Them1 in response to thermogenic stimuli, suggesting a new model for the regulation of energy expenditure within BAT. For this, Them1 activity is transiently silenced during peak thermogenesis by relocation away from its active site in puncta, which are located in close proximity to LD and mitochondria.

Our prior observations indicate that the activity of Them1 in brown adipocytes is to suppress the mitochondrial ß-oxidation of LD-derived fatty acids^11^. In keeping with this possibility, we observed that Them1 localized to puncta in the absence of adrenergic stimulation both in vitro and in vivo. These structures are juxtaposed with mitochondria and LD such that they could be expected to regulate fatty acid trafficking between these two organelles. Moreover, experiments in iBAs cells transduced with recombinant adenovirus revealed that full-length Them1 in unstimulated cells and the AAA-Them1 mutant, which is unresponsive to phosphorylation, were localized to puncta and were active in suppressing norepinephrine-stimulated oxygen consumption. These results are similar to our previous findings in cultured primary brown adipocytes, where full-length Them1 actively suppressed norepinephrine-stimulated oxygen consumption^11^, although puncta formation and dissolution were not investigated in that study.

By contrast, our data demonstrated that Them1 in its diffuse configuration was ineffective at suppressing oxygen consumption. This result was consistent with its relocation away from LD and mitochondria and suggests that S15 phosphorylation, and to a lesser degree S25 phosphorylation, both relocates and inactivates Them1. Our informatics, functional, and inhibitor data suggest that PKCβ is the kinase responsible for S15 phosphorylation. Interestingly, PKCβ appears to be involved in coordinating thermogenesis, as evidenced by increased rates of fatty acid oxidation in both BAT and white adipose tissue in mice globally lacking this kinase^15^.

The important role played by Them1-containing puncta in inhibiting the mitochondrial oxidation of LD-derived fatty acids begs the question about the composition of puncta, including how they are formed and why they disappear after phosphorylation. Our data using correlative light and electron microscopy suggest that Them1 constitutively resides in biomolecular condensates, also known as membraneless organelles, much like the nucleoleus, stress granules, and p-bodies^24,25^. Biomolecular condensates form by liquid-liquid phase separation or condensation^26^, which favors aggregation. The resulting structure often resembles an aggregation of round liquid droplets^21^, which is the result we found here for Them1 in puncta by electron microscopy. The driving force underlying the formation of biomolecular condensates is the exchange of macromolecular/macromolecular and water/water interactions under conditions that are energetically favorable^24^. What these conditions might be for Them1 are unknown but may be related to temperature, localized pH changes, or co-solutes that are expressed. Typical phase diagrams can define the conditions needed for condensation, which would be required to fully understand the localization of Them1 to puncta in vitro and in vivo.

One consistent signature of biomolecular condensates is the concept that their formation includes scaffold and client proteins^24^. Scaffold molecules are drivers of the condensation process and clients are molecules that partition into the condensate. It is not clear into which category Them1 fits, although the characteristics of its molecular structure suggest that it could be the primary scaffold protein. Phase separation of scaffold proteins and the subsequent partitioning of their clients are driven by network interactions for which two distinct classes of protein architecture are thought to be involved^24^. The first is multiple folded (SH3) domains that interact with short linear motifs^24^. Since Them1 does not have an SH3 domain, as determined by using the Simple Modular Architecture Research Tool web server^27^, this could not be the sequence element driving puncta formation. Second is intrinsically disordered regions^24^, for which Them1 has a strong candidate region near the amino terminus; amino acids 20-45 represents a highly disordered area that includes the S25 phosphorylation site and the N-terminal 20 amino acids are strongly predicted to be a disordered binding region that also includes phosphorylation sites at S15 and S18. Because the N-terminal 36 amino acids of Them1 are sufficient to drive puncta formation in iBAs cells, our data strongly supports that this region is involved in puncta formation and is the most likely area to investigate in the Them1 structure. Alternatively, Them1 may function as a client. Clients often have molecular properties similar to scaffolds including intrinsic disorder^25^, also making this role for Them1 a distinct possibility.

Dissolution of biomolecular condensates can be used to change the available concentration of cellular proteins to affect reaction efficiency^24^, and is likely important for Them1 to reduce its local concentration at LD and mitochondria subsequently increasing fatty acid availability for β-oxidation in mitochondria and increasing thermogenesis. The phosphorylation state of proteins regulates their ability to assemble in biomolecular condensates, with threonine phosphorylation resulting in condensation and serine phosphorylation reducing retention in a dose-dependent manner^21^. Phosphorylation events were shown to regulate phase separation, aggregation, and the retention of mRNA’s in FUS proteins, which are neuronal inclusions of aggregated RNA-binding protein fused in sarcoma^28,29^. It is thus phosphorylation at S15 and likely to some extent S25, which are highly conserved residues in Them1, that reduces its retention in puncta as evidenced in inhibitor studies and by genetic manipulation of Them 1. Norepinephrine-induced protein phosphorylation events during cold exposure in addition to a 3-fold increase in Them1 protein concentration over time^11^, likely creates equilibrium between active Them1 in puncta and inactive Them1 that is diffusely localized in the cytoplasm. Our data thus suggest that the delicate balance of Them1 protein concentration, its phosphorylation status, and environmental factors that promote Them1 phase transitions dictate the rate of thermogenesis in BAT.

Together, our results reveal an unexpected mechanism for the regulation of Them1, a protein that reduces fatty acid availability for β-oxidation in mitochondria and limits thermogenesis. Manipulation of Them1 localization in BAT may provide a new therapeutic target for the management of metabolic disorders including obesity and non-alcoholic fatty liver disease.

## Methods

### Culture of brown adipocytes

A brown adipose cell line (iBAs cells), which are immortalized preadipocytes from mouse brown adipose tissue^30^, were obtained from Dr. Bruce Spiegelman. iBAs cells were cultured in Dulbecco’s Modified Eagle’s medium (DMEM; Gibco, Grand Island, NY), 20% fetal bovine serum (FBS; Gibco), 1% penicillin/streptomycin solution (P/S; Gibco), and 2 μM HEPES (Gibco). Confluent cells were induced to differentiate from preadipocytes by adding induction cocktail containing 5μM dexamethasone (Sigma-Aldrich, St. Louis, MO), 125 μM indomethacin (Sigma-Aldrich), 0.02 μM insulin (Sigma-Aldrich), 500 μM IBMX (Life Technologies, Carlsbad, CA), 1 nM triiodothyronine (Sigma-Aldrich), and 1 μM rosiglitazone (Sigma-Aldrich) in medium consisting of DMEM, 10% FBS, and 1% P/S. After 48 h of induction, the medium was replaced by maintenance medium consisting of DMEM, 10% FBS, 1% P/S with 0.02 μM insulin, 1 nM triiodothyronine, and 1 μM rosiglitazone. The maintenance medium was replaced every two days. For immunostaining and confocal applications, the cells were plated on glass coverslips, which were pre-treated with 0.01% collagen type I (Sigma-Aldrich) in 0.01 M acetic acid solution.

### Production of DNA constructs

The full-length cDNA for Them1 (from Mus musculus:) was cloned and linked to EGFP plus an engineered ascorbate peroxidase (APEX2) fusion protein the C-terminus. The codons for S15, S18 and S25 were mutated to alanine (A) or aspartic acid (D) individually, in pairs, or in-all by using the QuikChange mutagenesis kit (Agilent, Santa Clara, CA). The Δ1-36 mutation includes deletion of the first 36 amino acids at the N-terminus of Them1, whereby Them 1 begins at amino acid 37 (M; AUG) in the gene sequence. The full-length cDNA for Them1 was cloned into a pVQAd CMV K-NpA vector (ViraQuest, North Liberty, IA) along with C-terminal EGFP downstream of the CMV promoter. The Them1 adenovirus was created by ViraQuest.

### Heterologous Them1 expression in cultured iBAs cells

Plasmid transfection with Lipofectamine 3000 (Thermo Fisher Scientific, Waltham, MA) was conducted 48 h after brown adipocytes were induced to differentiate. Purified DNA and P3000 reagent were mixed with Opti-MEM. The mixture was then added into Opti-MEM containing Lipofectamin 3000. The three reagents were allowed to incubate at room temperature for 10 min and then added to the cells. After 7 h of transfection, the medium was replaced by maintenance media. After 48 h, the cells were assayed. The same ratio of cells to DNA was used for each experiment but due to the very low transfection efficiency, it was not possible to assay Them1 concentration by Western blot.

For experiments using adenoviral vectors to express Them1, recombinant adenoviruses including Ad-Them1-EGFP, Ad-AAA-EGFP, Ad-DDD-EGFP, or Ad-EGFP alone were added to the medium for 24 h after the first day of differentiation. Adenoviral vectors were initially screened for MOI versus cell protein expression using our Them1 antibody^11^, mRNA expression using quantitative RT-PCR (Supplementary Fig. 2b,c), and cell viability using the crystal violet assay^31^, which was nearly 100% with all MOI’s used from 10 through 80 (not shown). An MOI of 40 was used for all studies because the intracellular Them1 concentration was most similar to that induced in BAT by cold exposure (Supplementary Fig. 2b).

### Fixation and Oil Red O staining for LD

Differentiated iBAs cells were washed with phosphate buffered saline (PBS) and then fixed with 4% paraformaldehyde in PBS for 20 min at room temperature. Cells were washed twice before staining with 0.3% Oil Red O solution in 60% isopropanol (Sigma Aldrich) at room temperature. The cells were washed with PBS and then mounted with ProLong Diamond Antifade Mountant with DAPI (ThermoFisher) to identify nuclei.

### Immunostaining for LD, mitochondria, and plasmid-expressed Them1

Differentiated iBAs cells were fixed, as above, and then stained for perilipin (anti-Plin1) to identify LD, Tomm20 (anti-Tomm20) to detect mitochondria, or anti-EGFP to visualize the plasmid-transfected Them1-EGFP. For staining Them1 in tissue sections of BAT, antigen retrieval was done in paraffin sections with citrate buffer, pH 6.0, and then the sections were stained with anti-Them1 antibody, anti-Plin1, and DAPI to identify nuclei. Details of the antibodies: anti-Them1^11^, rabbit, 1:100 for immunostaining and 1:1000 for Western blots; anti-Tomm20, Santa Cruz Biotech (Dallas TX, USA), rabbit, sc-11415, 1:200; anti-Plin1, Fitzgerald (Acton MA, USA), guinea pig, 20R-PP004,1:500; anti-EGFP, Novus Biologicals (Littleton CO, USA), goat, NB100-1678, 1:500; anti-actin, Abcam (Cambridge, MA, USA), mouse, Ab3280,1:1000. Secondary antibodies labeled with HRP, AlexaFluor 488, 647, or Cy3 were purchased from Jackson ImmunoResearch (West Grove PA, USA).

### Confocal Microscopy

The localization of fluorescence signal in cultured iBAs cells was evaluated using a Zeiss LSM880 confocal microscope system with or without Airyscan. Images were acquired at high resolution as 2 µm z-stack slices through the thickness of cells and assembled in 3-D using Volocity image processing software.

For confocal imaging of living cells, a time-lapse image was created to capture the dynamics of protein expression and localization, signal transduction, and enzyme activity. Cells grown on coverslips were placed in the incubator chamber of a Zeiss LSM880 inverted confocal microscope system to visualize EGFP expression (linked to Them1). The interior of the chamber was kept at a constant 37° C, and set to a 5% CO_2_ level and a 95% O_2_ level. An image was captured at T0, and immediately following the application of a treatment. The microscope then automatically tracked, focused, and photographed cells every 30 min for 4-5 h. These images are also taken in the high resolution z-stack format and analyzed using Volocity software.

### Treatment of iBAs cells to explore metabolic pathways

To explore the metabolic pathway leading to re-organization of intracellular Them1, various treatments were applied to cultured brown adipocytes as follows. Norepinephrine (Sigma Aldrich; St. Louis, MO), a neurotransmitter, was used to mimic cold exposure. Forskolin (FSK) (Selleckchem, Houston, TX), a membrane permeable labdane diterpene produced from the *Coleus* plant, was used to activate adenylyl cyclase. This treatment, through the activation of cAMP, activates PKA. PMA (LC Laboratories, Woburn, MA) was used to activate PKC. NE, adenylyl cyclase, and PKC were selected for activation as they represent three key control points in the pathway allowing the characterization of all upstream regulators. PKI ([14-22], Myristoylated) (Invitrogen, Camarillo, CA), a synthetic peptide inhibitor of PKA, was added to inhibit PKA activation. Atglistatin (Selleckchem, Houston, TX) was used to selectively inhibit the processing of triacylglycerol to fatty acids via the formation of diacylglycerol by ATGL. Ruboxistaurin, or LY333531 (Selleckchem, Houston, TX), an isozyme-selective inhibitor of PKC that competitively and reversibly inhibits PKC-βI and PKC-βII, was added to inhibit PKCβ activity.

### Correlative light and electron microscopy of Them1 in iBAs cells

Differentiated iBAs cells transfected with a plasmid containing Them1 linked to APEX2 were fixed and the APEX2 developed using an ImPACT peroxidase substrate kit (Vector Labs, Burlingame, CA). Cells were then frozen using a Wohlwend Compact 02 High Pressure Freezer. Frozen cells underwent super-quick freeze substitution and embedding using the protocol established by McDonald^32^. In 0.5 micrometer thick sections, positive cells stained brown by light microscopy. If positive cells were identified, four consecutive levels of serial ultra-thin sections were obtained with one thick section for light microscopy between levels. Light microscopy images, taken with an Axioimager (Zeiss) equipped with a color ccd camera were used to follow puncta in the sections and to determine the position of positive cells in ultra-thin sections. Images were taken on a JEOL 1400 electron microscope equipped with a Gatan (Gatan, Pleasanton, CA) Orius SC1000 camera.

### O_2_ consumption rates in iBAs cells

Oxygen consumption rates (OCR) were measured in iBAs cells using a Seahorse XFe96 (Agilent Technologies, Santa Clara, CA, USA). Briefly, iBAs cells were seeded at a density of 1,000 cells/well in collagen-coated Seahorse XF96 cell culture microplates two days before induction. On day 1 after induction, iBAs cells were incubated with Ad-Them1-EGFP, Ad-AAA-EGFP, Ad-DDD-EGFP, or Ad-EGFP alone. One day after adenovirus infection, the medium was changed to the maintenance medium supplemented with 1μM NE. After two days, CO_2_ was withdrawn from the cultures for 1 h at 37°C in Krebs-Henseleit buffer (pH 7.4) containing 2.5 mM glucose, 111 mM NaCl, 4.7 mM KCl, 2 mM MgSO_4_-7H_2_O, 1.2 mM Na_2_HPO_4_, 5 mM HEPES and 0.5 mM carnitine (denoted as KHB). Basal oxygen consumption rate was measured in in KHB for 18 min and then 10 μM NE was injected through the Seahorse injection ports for a final concentration of 1 μM in each well. NE responses were measured for 54 min after injections as previously described^11^.

After experiments, cell viability/cell number was determined using the NucRed™ Live 647 ReadyProbes™ assay (Thermo Fisher Scientific, Waltham, MA, USA) and the color read using a Spectramax i3x (Molecular Devices, San Jose, CA, USA). Cell viability was calculated as the percent of viable cells transfected with EGFP alone compared to the other conditions. All values for OCR were normalized to cell number.

### Activation of BAT in mice, in vivo

Male C57BL/6J mice (7 weeks) were purchased from the Jackson Laboratory (Bar Harbor, ME) and housed in an AAALAC International Facility at Weill Cornell Medical College. Animal use and euthanasia protocols were approved by the Institutional Animal Care and Use Committee at Weill Cornell Medical College.

Mice were placed in a Promethion metabolic cage (Sable Systems International, Las Vegas, NV) with one mouse/cage. Mice were housed at 22° C for 24 h and then moved to 30° C for experiments. Room temperature, or 22° C, is below the thermoneutral zone of 30° C for mice, which activates Them1 expression in BAT by mild cold exposure^11^. Experimental mice were injected subcutaneously with one dose of CL316,243 (CL, Sigma-Aldrich, 1 mg/kg body weight) in PBS, which is a β3 adrenergic receptor agonist. Control mice were injected with PBS alone. Subscapular BAT was excised from mice after euthanasia at T0, 1, 2, or 4 h after injection. Tissues were fixed overnight in 10% neutral buffered formalin and then processed. Paraffin sections were used for H&E stained samples and for immunocytochemistry as described above.

### Image processing and quantification of fluorescence signal

For experiments aimed to quantify changes in intercellular Them1 expression in iBAs cells, the volume of Them1, size, and/or distribution of puncta was quantified in the 3-D volume using Volocity software (Quorum Technologies, Ontario, Canada) or Imaris software (Oxford Instruments, Concord, MA, USA). The strategy used for both iBAs cells and for tissue Them1 expression was that puncta represent concentrated Them1 that occupies a small cellular volume whereas diffuse Them1 is distributed evenly throughout the entire cell volume resulting in a large cellular volume. Them1 fluorescence signal, which represented all pixel intensities, was measured by using a normalized histogram for the distribution of Them1 protein at each 12-bit (0 - 4096) intensity value. This strategy does not stratify the signal into intense (as found in puncta) or weak (as found in the diffuse state), but rather identifies the volume of total intracellular Them1 fluorescence irrespective of intensity. The sum was then normalized to a measurement of total cell volume and graphed using SigmaPlot software (SystatSoftware, San Jose, CA). Intensity measurements, size, and distribution measurements for each spot from 0-4096 in the 3-D volume were quantified using Imaris software.

### Bioinformatics

The amino acid sequence of Them1 from *Mus musculus* was acquired from UniProt^33^ and analyzed with multiple servers. The IUPRED2A server was used to predict intrinsic disorder and ANCHOR binding regions^34–37^. The Prion-like Amino Acid Composition (PLAAC) server was used to predict prion like regions through the prion like domains (PLD) and FOLD curves^38^. The propensity of Them1 to phase separate through long-range planar pi-pi contacts was predicted using a server generated by the laboratory of Professor Julie Forman-Kay (PScore)^39^. The Motif Scan server (https://myhits.isb-sib.ch/cgi-bin/motif_scan) was utilized to examine the first 20 amino acids for known motifs^23^.

### Statistics and reproducibility

Data were analyzed using SigmaPlot software with a *P* value less than 0.05 considered significant. For comparison of data from many groups, one-way analysis of variance test was used to determine statistical significance. If variances were not normal, non-parametric ANOVA was done using ranks. For experiments with multiple times and treatments, 2-way analysis was done to determine significance between groups. Error bars indicate s.e.m. in all experiments. All data are from 3 or more independent experiments with results obtained from at least 3 replicates within each group.

## Data Availability

The authors declare that the main data supporting the findings of this study are available within the article and its Supplementary Information files. Adenoviral vectors or plasmid constructs are available upon reasonable request to the corresponding authors.

## Acknowledgements

The authors thank Dr. Bruce Spiegelman, Dana-Farber Cancer Institute, Harvard Medical School for providing the iBAs cells, Mr. Kyle Smith for his technical help with electron microscopy, and Mr. Aniket Gad for expert advice on image processing and for providing the data in Figure 2. This work was supported by the National Institutes of Health (RO1 DK 103046 to D.E.C., S.J.H, and E.O.), the Harvard Digestive Diseases Center (P30 DK034854 to S.J.H.), and the National Institutes of Health shared-instrumentation grant program for the High Pressure Freezer (S10 OD019988-01 to S. J.H.). H.T.N is the recipient of a Pinnacle Research Award from the AASLD Foundation and T.I.K. is the recipient of a Weill Cornell Department of Medicine Pre-Career Award and acknowledges support from NIH T32DK007533.

## Author Contributions

Y.L. did the cell culture experiments; N.I. did the Seahorse work; A.G. assisted with pathway experiments and image processing; H.T.N. and T.I.K. performed the animal experiments; M.B. assisted with experiments; L.H.A. did the immunostaining; M.T. and E.O. did the Them1 bioinformatic analysis; S.J.H. did CLEM, co-mentored Y.L., assisted with experimental design, and wrote the paper. D.E.C. co-mentored Y.L., assisted with experimental design, and edited the paper. All authors reviewed and edited the final manuscript.

## Materials & Correspondence

Correspondence and requests for materials should be addressed to S.J.H. or D.E.C.

## References

1. Benador, I. Y., Veliova, M., Mahdaviani, K., Petcherski, A., Wikstrom, J. D., et al. Mitochondria bound to lipid droplets have unique bioenergetics, composition, and dynamics that support lipid droplet expansion. Cell Metab. 27, 869–885 (2018).

2. Rambold, A. S., Cohen, S., & Lippincott-Schwartz, J. Fatty acid trafficking in starved cells: regulation by lipid droplet lipolysis, autophagy, and mitochondrial fusion dynamics. Dev. Cell 32, 678–692 (2015).

3. Cannon, B. & Nedergaard, J. Brown adipose tissue: function and physiological significance. Physiol. Rev. 84, 277–359 (2004).

4. Deiuliis, J. A., Liu, L. F., Belury, M. A., Rim, J. S., Shin, S., & Lee, K. β_3_-adrenergic signaling acutely down regulates adipose triglyceride lipase in brown adipocytes. Lipids 45, 479–489 (2010).

5. Brasaemle, D. L. Thematic review series: adipocyte biology. The perilipin family of structural lipid droplet proteins: stabilization of lipid droplets and control of lipolysis. J. Lipid Res. 48, 2547–2559 (2007).

6. Ellis, J. M., Li, L. O., Wu, P. C., Koves, T. R., Ilkayeva, O., et al. Adipose acyl-CoA synthetase-1 directs fatty acids toward beta-oxidation and is required for cold thermogenesis. Cell Metab 12, 53–64 (2010).

7. Grevengoed, T. J., Klett, E. L., & Coleman, R. A. Acyl-CoA metabolism and partitioning. Annu. Rev. Nutr. 34, 1–30 (2014).

8. Tillander, V., Alexson, S. E. H., & Cohen, D. E. Deactivating fatty acids: acyl-CoA thioesterase-mediated control of lipid metabolism. Trends Endocrinol. Metab. 28, 473–484 (2017).

9. Adams, S. H., Chui, C., Schilbach, S. L., Yu, X. X., Goddard, A. D., et al. BFIT, a unique acyl-CoA thioesterase induced in thermogenic brown adipose tissue: cloning, organization of the human gene and assessment of a potential link to obesity. Biochem. J. 360, 135–142 (2001).

10. Zhang, Y., Li, Y., Niepel, M. W., Kawano, Y., Han, S., et al. Targeted deletion of thioesterase superfamily member 1 promotes energy expenditure and protects against obesity and insulin resistance. Proc. Natl. Acad. Sci. U. S. A 109, 5417–5422 (2012).

11. Okada, K., LeClair, K. B., Zhang, Y., Li, Y., Ozdemir, C., et al. Thioesterase superfamily member 1 suppresses cold thermogenesis by limiting the oxidation of lipid droplet-derived fatty acids in brown adipose tissue. Mol. Metab. 5, 340–351 (2016).

12. Huttlin, E. L., Jedrychowski, M. P., Elias, J. E., Goswami, T., Rad, R., et al. A tissue-specific atlas of mouse protein phosphorylation and expression. Cell 143, 1174–1189 (2010).

13. Horn, H., Schoof, E. M., Kim, J., Robin, X., Miller, M. L., et al. KinomeXplorer: an integrated platform for kinome biology studies. Nat. Methods 11, 603–604 (2014).

14. Blom, N., Sicheritz-Ponten, T., Gupta, R., Gammeltoft, S., & Brunak, S. Prediction of post-translational glycosylation and phosphorylation of proteins from the amino acid sequence. Proteomics 4, 1633–1649 (2004).

15. Bansode, R. R., Huang, W., Roy, S. K., Mehta, M., & Mehta, K. D. Protein kinase C deficiency increases fatty acid oxidation and reduces fat storage. J. Biol. Chem. 283, 231–236 (2008).

16. Mehta, K. D. Emerging role of protein kinase C in energy homeostasis: a brief overview. World J. Diabetes 5, 385–392 (2014).

17. Huang, W., Bansode, R., Mehta, M., & Mehta, K. D. Loss of protein kinase Cβ function protects mice against diet-induced obesity and development of hepatic steatosis and insulin resistance. Hepatology 49, 1525–1536 (2009).

18. Krisko, T. I., Nicholls, H. T., Bare, C. J., Holman, C. D., Putzel, G. G., et al. Thermogenesis from glucose homeostasis in microbiome-deficient mice. Cell Metab. in press, (2020).

19. Song, A., Dai, W., Jang, M. J., Medrano, L., Li, Z., et al. Low- and high-thermogenic brown adipocyte subpopulations coexist in murine adipose tissue. J. Clin. Invest. 130, 247–257 (2020).

20. Friend, D. S. & Brassil, G. E. Osmium staining of endoplasmic reticulum and mitochondria in the rat adrenal cortex. J. Cell Biol. 46, 252–266 (1970).

21. Li, P., Banjade, S., Cheng, H.-C., Kim, S., Chen, B., et al. Phase transitions in the assembly of multivalent signaling proteins. Nature 483, 336–340 (2012).

22. Pak, C. W., Kosno, M., Holehouse, A. S., Padrick, S. B., Mittal, A., et al. Sequence Determinants of Intracellular Phase Separation by Complex Coacervation of a Disordered Protein. Mol. Cell 63, 72–85 (2016).

23. Sigrist, C. J., Cerutti, L., de, C. E., Langendijk-Genevaux, P. S., Bulliard, V., Bairoch, A., & Hulo, N. PROSITE, a protein domain database for functional characterization and annotation. Nucleic Acids Res. 38, D161–D166 (2010).

24. Alberti, S., Gladfelter, A., & Mittag, T. Considerations and challenges in studying liquid-liquid phase separation and biomolecular condensates. Cell 176, 419–434 (2019).

25. van Leeuwen, W. & Rabouille, C. Cellular stress leads to the formation of membraneless stress assemblies in eukaryotic cells. Traffic 20, 623–638 (2019).

26. Martin, E. W. & Mittag, T. Relationship of sequence and phase separation in protein low-complexity regions. Biochemistry 57, 2478–2487 (2018).

27. Letunic, I. & Bork, P. 20 years of the SMART protein domain annotation resource. Nucleic Acids Res. 46, D493–D496 (2018).

28. Han, T. W., Kato, M., Xie, S., Wu, L. C., Mirzaei, H., et al. Cell-free formation of RNA granules: bound RNAs identify features and components of cellular assemblies. Cell 149, 768–779 (2012).

29. Monahan, Z., Ryan, V. H., Janke, A. M., Burke, K. A., Rhoads, S. N., et al. Phosphorylation of the FUS low-complexity domain disrupts phase separation, aggregation, and toxicity. EMBO J. 36, 2951–2962 (2017).

30. Uldry, M., Yang, W., St-Pierre, J., Lin, J., Seale, P., & Spiegelman, B. M. Complementary action of the PGC-1 coactivators in mitochondrial biogenesis and brown fat differentiation. Cell Metab. 3, 333–341 (2006).

31. Seo, J. H., Fox, J. G., Peek, R. M., & Hagen, S. J. *N*-methyl D-aspartate channels link ammonia and epithelial cell death mechanisms in *Helicobacter pylori* infection. Gastroenterology 141, 2064–2075 (2011).

32. McDonald, K. L. Rapid embedding methods into epoxy and LR White resins for morphological and immunological analysis of cryofixed biological specimens. Microsc. Microanal. 20, 152–163 (2014).

33. UniProt: a worldwide hub of protein knowledge. Nucleic Acids Res. 47, D506–D515 (2019).

34. Meszaros, B., Erdos, G., & Dosztanyi, Z. IUPred2A: context-dependent prediction of protein disorder as a function of redox state and protein binding. Nucleic Acids Res. 46, W329–W337 (2018).

35. Dosztanyi, Z. Prediction of protein disorder based on IUPred. Protein Sci. 27, 331–340 (2018).

36. Dosztanyi, Z., Csizmok, V., Tompa, P., & Simon, I. The pairwise energy content estimated from amino acid composition discriminates between folded and intrinsically unstructured proteins. J. Mol. Biol. 347, 827–839 (2005).

37. Meszaros, B., Simon, I., & Dosztanyi, Z. Prediction of protein binding regions in disordered proteins. PLoS Comput. Biol. 5, e1000376 (2009).

38. Lancaster, A. K., Nutter-Upham, A., Lindquist, S., & King, O. D. PLAAC: a web and command-line application to identify proteins with prion-like amino acid composition. Bioinformatics. 30, 2501–2502 (2014).

39. Vernon, R. M., Chong, P. A., Tsang, B., Kim, T. H., Bah, A., et al. Pi-Pi contacts are an overlooked protein feature relevant to phase separation. Elife. 7, (2018).

