## Supplementary Figures for "Thioesterase Superfamily Member 1 Undergoes Stimulus-coupled Reorganization to Regulate Metabolism"

```
                                15 18      25
Human            1 MIQNVGNHLRRGLASVFSNRTSRKSALRAG--N-D-SAMADGEG 40
Gorilla          1 MIQNVGNHLRRGLASVFSNRTSRKSALRAG--N-D-SAMADGEG 40
Flying Lemur     1 MIQNVGSHLRRGFASVFSSRTSRKSASRAG--D-NDFAMAEGEG 41
Wolf             1 MIQNVGNHLRRGIASVFSSRTSRKSASRAE--K-DSGAMADGEG 41
Sea Lion         1 MIQNVGNHLRRGIASVFSSRTSRKSASRSE--H-ADGAMADGEG 41
Fruit Bat        1 MIQNVGNHLRRSFASMFSSRQSRKSTSRAE--D-D-GAMADGEG 40
European Rabbit  1 MIQTVGSHLRRGFASVFSSRTSRKSASRAG--D-ADGAMADGEG 41
Camel            1 MIQNVGNHLRRGLASVFSSRTSRKSALRAE--N-T---MAEGEG 38
Alpaca           1 MIQNVGNHLRRGLASVFSSRTSRKSASRAE--N-T---MAEGEG 38
Minke Whale      1 MIQNVGNHLRRGLASVFSNRASRKSASHTG--N-N---MAEGEG 38
Baiji            1 MIQNVGNHLRRGLASVFSNRASRKSASRTE--N-N--NMAEDEG 37
Artic Squirrl    1 MIQNVGNHLRRGLASVFSSRTSRKSVSRAG--D-DN-DMEEGEG 40
Cattle           1 MIQTVGNHLRRGLASVFSNRTSRKSASRTD--S-D--NMADGEG 39
White Rhino      1 MIQNVGTHLRRGLASVFSSRTSRKSASRAE--K-DGGVMAEGEG 41
Meerkat          1 MIQNVGNHLRRSIASVFSSRTSRKSASRAE--K-D-GAMADGEG 40
Sheep            1 MIQTVGSHLRRGLASVFSNRTSRKSASRTD--S-D--NMADGEG 39
Cheetah          1 MIQNVGNHLRRGFASVFSSRLSRKSASHAE--K-DDGAMAGGEG 41
Goat             1 MIQTVGSHLRRGLASVFSNRTSRKSASRTD--S-D--NMADGEG 39
Beluga Whale     1 MIQNVGNHLRRGLASVFSNRASRKSASRTE--N-N--NMAEDKG 39
Wild Boar        1 ---------MQGLASVFSNRASKKSAPRSE--N----NMGDGEG 26
Cat              1 MIQNVGNHLRRGFTSVFSSRLSRKSASHAE--K-DDGAMAGGEG 41
Leopard          1 MIQNVGNHLRRGFASVFSSRMSRKSASHAE--K-DDGAMAGGEG 41
Horse            1 MIQNVGNHLRRGLSSVFSSRSARKSASRAD--K-DGGAMAAGEG 41
Naked Mole Rat   1 MIQNVGSHLRRSFASVFSNRTSRKSVSRAG--D-ED-TMANSEG 40
Armadillo        1 MIQNVGNHLRRGFTSVFSGRTSRKSASRARAED-ADGAMAD-EG 42
Mouse            1 MIQNVGNHLRRGFASMFSNRTSRKSISHPE--SGDPPTMAEGEG 42
Wombat           1 -MQNIGNQLRRGLTSIFSGRASRKQAAQGR--D-PAGTM---ET 37
```

**Supplementary Fig. 1** Conservation of the *Them1* mRNA sequence for amino acids S15, S18, and S25 at the N-terminus between human, mouse, and other species.

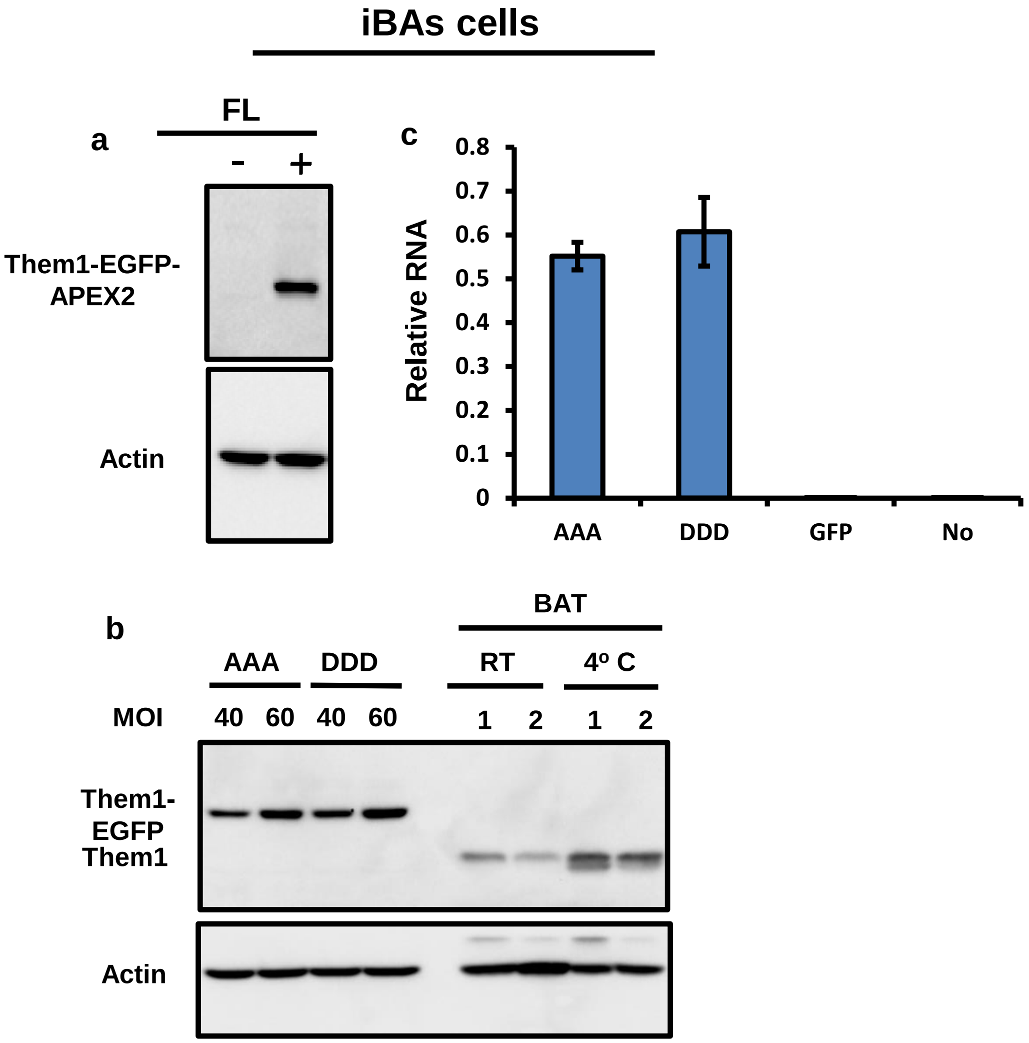


**Supplementary Fig. 2 Expression levels of Them1 in immortalized brown adipose (iBAs) cells and in brown adipose tissue (BAT). a,** iBAs cells do not constitutively express Them1 protein in culture. Transfection of a Them1-EGFP/APEX2 plasmid results in Them1 expression. Data are representative of *n* = 2 independent experiments. **b,** Plasmids expressing the AAA- or DDD-Them1 mutant in iBAs cells at an MOI of 40 express Them1 to a similar level as BAT tissues from mice exposed to cold (4º C) for 48 h. Data are representative of *n* = 3 independent experiments. For BAT tissue, cold exposure for 48 h increases Them1 protein expression. Data are from 2 mice (1, 2) for room temperature (RT) and from 2 mice (1,2) exposed to the cold (4º C) for 48 h. **c,** Results from RT-PCR to demonstrate relative mRNA expression in iBAs cells transfected with the AAA- or DDD-Them1 mutant plasmid compared to cells transfected with GFP alone or cells transfected with an empty vector containing no Them1. Data are representative of *n* = 3 independent experiments. Western blot data were not processed or modified from the original scans.

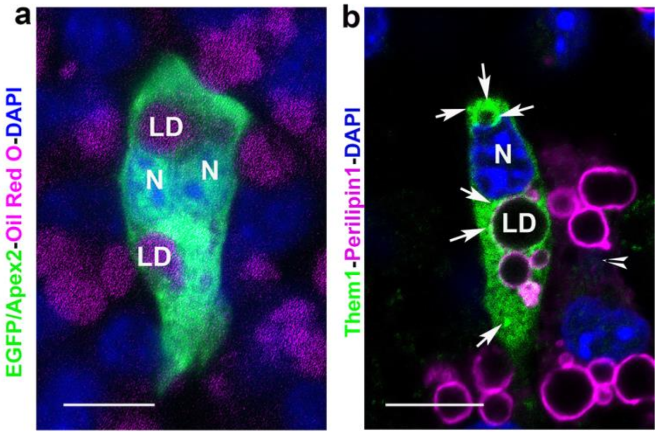


**Supplementary Fig. 3. EGFP, *per se*, does not cause puncta formation in iBAs cells transfected with Them1-EGFP/Apex2. a,** iBAs cells were transfected with a plasmid containing EGFP and Apex2 alone, without Them1. Note that the EGFP fluorescence signal is distributed uniformly throughout the cell cytoplasm and nuclei (N) excluding lipid droplets (LD). Data are representative of *n* = 2 independent experiments. **b,** iBAs cells were transfected with Them1 containing no EGFP/Apex2. Note that Them1 formed puncta (arrows) in the cytoplasm and near lipid droplets (LD). Data are representative of *n* = 3 cells from one experiment. N, nuclei. Scale bars, 10 μm.

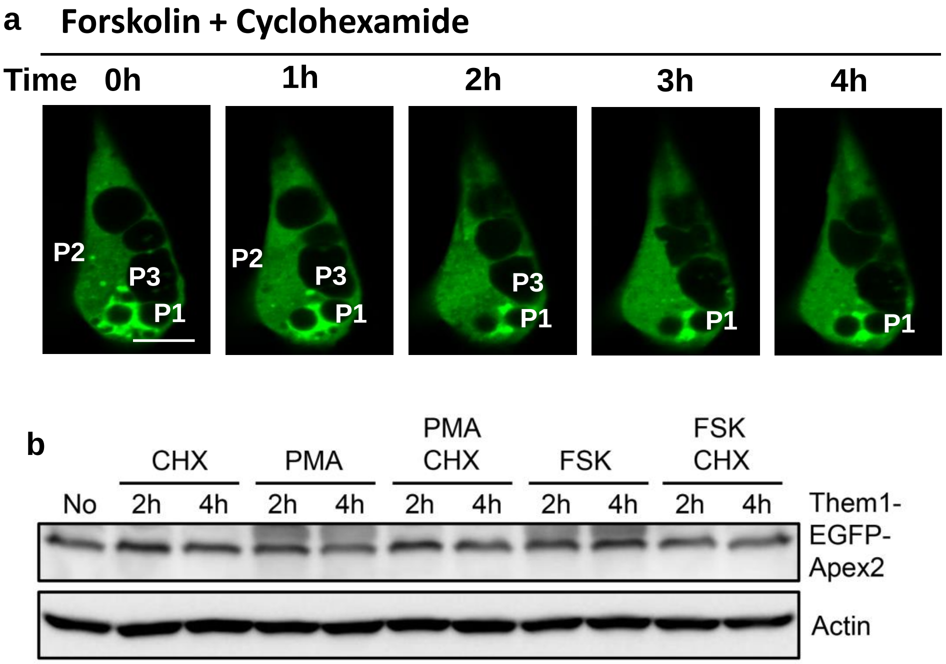


**Supplementary Fig. 4 Blocking protein synthesis with cyclohexamide has no effect on the diffusion of Them1 from puncta or the expression of intracellular Them1 in iBAs cells after stimulation with forskolin or PMA. a,** Cyclohexamide (0.36 mM) was added with forskolin to stimulate PKC activation and the dissolution of puncta (P). Numerous puncta, P1, P2, and P3, can be followed in the cell over time. Scale bar, 10 μm. **b**, Western blot from iBAs cells treated with cyclohexamide alone (CHX, 0.36 mM), PMA alone, PMA plus cyclohexamide, forskolin (FSK), or forskolin plus cyclohexamide. Data are representative of *n* = 3 independent experiments. Western blot data were not processed or modified from the original scans.

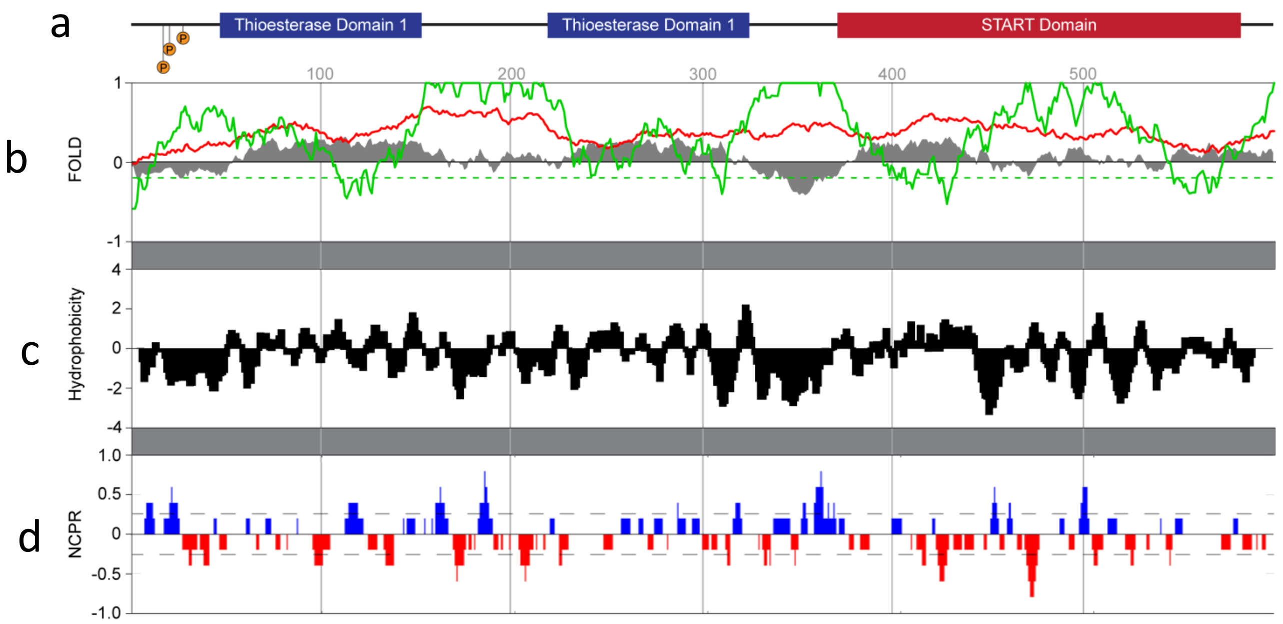


**Supplementary Fig. 5 Bioinformatic support for the analysis in Fig. 6. a,** Schematic diagram of the Them1 amino acid sequence aligned with the amino acid number from the N- to carboxy-terminus. **b,** Results from the FOLD server, which predicts regions of intrinsic disorder by PLAAC (green) and the Prion Aggregation Prediction Algorithm (PAPA-red). The FOLD index is in grey. This analysis confirms the PLD server analysis (Fig. 6c) that Them1 has no prion-like regions in the sequence. **c,** Hydrophobicity analysis shows that the Them1 sequence contains a high proportion of charged residues with a patch of basic residues followed by a patch of acidic residues. These patches could potentially engage in non-covalent crosslinking with other proteins or itself to drive a phase transition. **d,** NCPR shows the net charge per residue. This information can help to understand the loss of solubility of Them1, particularly in disordered regions. The dotted line indicates the significance threshold.
